# Differential expression of BMP antagonists, *gremlin* and *noggin* in hydra: antagonism between Wnt and BMP pathways

**DOI:** 10.1101/510602

**Authors:** Krishnapati Lakshmi Surekha, Samiksha Khade, Diptee Trimbake, Rohan Patwardhan, Siva Kumar Nadimpalli, Surendra Ghaskadbi

## Abstract

Mechanisms regulating BMP and Wnt signaling pathways have been widely studied in many organisms. One of the mechanisms by which these pathways are regulated is by binding of extracellular ligands. In the present study, we report studies with two BMP antagonists, *gremlin* and *noggin* from *Hydra vulgaris* Ind-Pune and demonstrate antagonistic relationship between BMP and Wnt pathways. *Gremlin* was ubiquitously expressed from the body column to head region except in the basal disc and hypostome. During budding, *gremlin* was expressed predominantly in the budding region suggesting a possible role in budding; this was confirmed in polyps with different stages of buds. *Noggin*, on the other hand, was predominantly expressed in the endoderm of hypostome, base of the tentacles, lower body column and at the basal disc in whole polyps. During budding, *noggin* was expressed at the sites of emergence of tentacles suggesting a role in tentacle formation. This was confirmed in alsterpaullone-treated polyps, which showed *noggin* expression as distinct spots where ectopic organizers and ectopic tentacles eventually formed. Using RT-PCR, we found that up-regulation of *Wnt* is accompanied with down-regulation of *BMP5-8b* demonstrating antagonism between the two pathways. Down-regulation of *noggin* and *gremlin*, however, occurred only after 24 h recovery. The data suggest that inhibition of BMP pathway by Wnt signaling in hydra does not directly involve *noggin* and *gremlin*. Our findings indicate that the BMP/Noggin antagonism evolved early for setting up and/or maintaining the head organizer while involvement of these BMP antagonists during vertebrate axial patterning are recent evolutionary acquisitions.

**Summary statement:** We show that setting up of the Organizer by BMP/Noggin antagonism and role of BMP inhibitors in tissue patterning are evolutionarily ancient, probably arising for the first time in hydra

## Introduction

Bone morphogenetic protein family, a subfamily of transforming growth factor-β (TGF-β) superfamily was originally identified for role of some of its members in formation and regeneration of bone *in vivo*. Subsequently, their roles in crucial developmental processes, such as, proliferation, migration, differentiation and fate specification of embryonic cells, morphogenesis and dorso-ventral patterning were discovered (Massagué, 1998; Komiya and Habas, 2008). Non-optimal BMP signaling leads to a variety of developmental abnormalities and disease conditions including cardiovascular diseases, symphalangism, diabetic nephropathy and several types of cancers (Kattamuri et al., 2017). BMPs initiate downstream signaling by two mechanisms. The first is a canonical smad-dependent pathway which involves phosphorylation of Serine/Threonine kinase receptors, Type I and Type II receptor complexes (BMPRI and BMPRII), and activation of smad-1/5/8. The second is a smad-independent pathway, which involves TGF-β Activated Tyrosine Kinase 1 (TAK1) and Mitogen Activated Protein Kinase (MAPK) (Zhang and Li, 2005). Similar to BMP pathway, Wnt family of secreted glycoproteins also act as powerful regulators of embryonic development, cell proliferation, migration, differentiation and cell fate specification. Extracellular glycosylated Wnt ligands bind to Frizzled receptor and co-receptors, such as, lipoprotein receptor-related protein (LRP) 5/6-, Ryk and Ror. Once initiated, signal transduction occurs through two distinct mechanisms: canonical or β-catenin-dependent pathway and a non-canonical β-catenin-independent pathway.

Mechanisms regulating BMP and Wnt signaling pathways and the regulatory networks involved in the downstream signaling pathways have been studied extensively in many organisms (Azpiazu et al., 1996; Carmena et al., 1998; Hoppler and Moon, 1998; Jin et al., 2001). Both the pathways can function either independently or exhibit overlapping expression patterns suggesting a crosstalk between them (Itasaki and Hoppler, 2009). One of the important mechanisms through which these two pathways interact is by secreted molecules that bind to the extracellular components of these pathways. Several molecules like Noggin, Chordin, Gremlin and Follistatin, which inhibit BMP signaling by binding to and neutralizing BMP ligands have been identified. Similarly, Cerberus and Connective tissue growth factor bind to Wnt and BMP, affecting both pathways (Bouwmeester et al., 1996). Though BMP pathway was initially thought to have originated along with the dorso-ventral axis in bilaterians (Finnerty, 2003), occurrence of components of this pathway in lower invertebrates like Cnidarians has raised several questions regarding their evolutionary functions in regulating axial patterning.

Hydra, a Cnidarian, has been a favorite model for studying morphogenesis and pattern formation because of its unique features, such as, maintenance of axial polarity, spectacular ability of regeneration, absence of cellular senescence, maintenance of stemness, and peculiar tissue dynamics. Several components of BMP and Wnt pathways have been characterized in hydra. Eleven Wnt ligands and components of the pathway, β-catenin and Tcf have been reported from hydra and their roles in the formation of head Organizer, axial patterning and tissue morphogenesis as well as in regeneration have been established (Hobmayer et al., 1996; 2001; Broun et al., 2005; Philipp et al., 2009). Similarly, role of *HyBMP5-8b*, a BMP5-8 orthologue, in tentacle formation and patterning the body axis has been reported (Reinhardt et al., 2004). *HySmad1*, a smad gene involved in nematocyte differentiation and oogenesis (Hobmayer et al., 2001), and inhibitors of BMP ligand, Chordin (Rentzsch et al., 2006) and Noggin (Chandramore et al., 2010) from hydra have also been reported. Our earlier work has shown that hydra Noggin induces secondary axis in *Xenopus* embryos and partially rescues UV-induced ventralization in *Xenopus* embryos through inhibition of BMP signaling, confirming its functional conservation in vertebrate embryos (Chandramore et al., 2010). Occurrence of components of BMP pathway in hydra thus point toward their origin before the divergence of Cnidarians and bilaterians (Reinhardt et al., 2004; Rentzsch et al., 2006). Though genes involved in axial patterning, such as, *Wnt, BMPs* and those encoding BMP inhibitors have been reported from hydra, their detailed structural and functional characterization remains to be done. In the present study, we report *gremlin*, a secreted BMP antagonist belonging to the DAN (<u>D</u>ifferential screening-selected gene <u>A</u>berrative in <u>N</u>euroblastoma) family for the first time and compare it to *noggin*, another BMP antagonist from *Hydra vulgaris* Ind Pune. We find that *gremlin* and *noggin* are differentially expressed in hydra suggesting roles in different biological processes. Further, we demonstrate an antagonistic relationship between Wnt and BMP pathways in hydra that does not directly involve *noggin* and *gremlin*.

## Results

### Identification of *gremlin* like gene from Hydra

Search for hydra *gremlin* sequence in the NCBI database led to identification of a predicted *gremlin-like 1* mRNA sequence (gi|449664166) from *Hydra magnipapillata*. Complete coding sequence of 483 bp amplicon was amplified from *Hydra vulgaris* Ind-Pune using primers designed based on the available predicted sequence (Fig.1a). The amplicon was cloned, sequenced and submitted to the Genbank with accession number KJ672500.

**Fig. 1.**
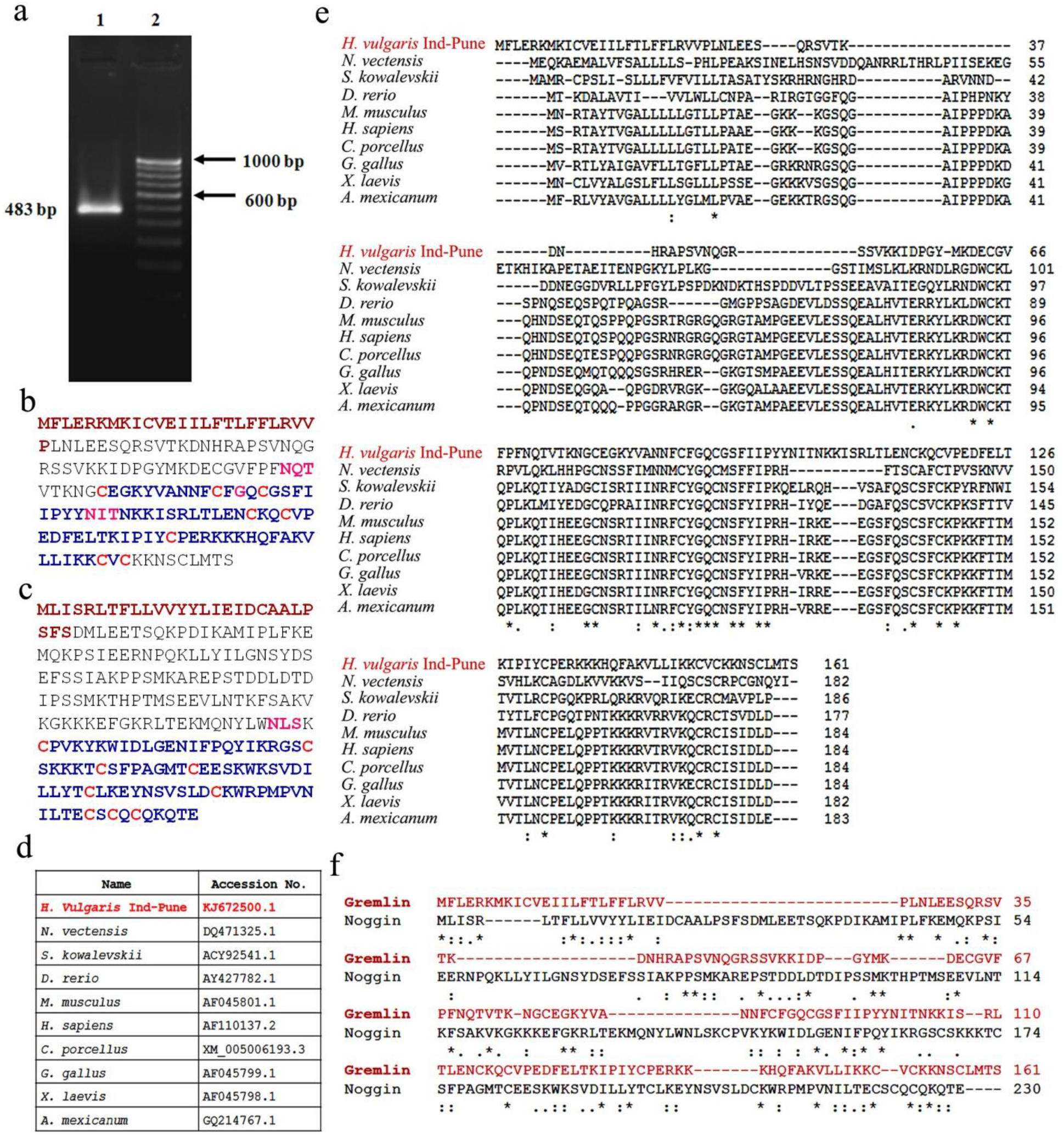
Identification of *gremlin* in hydra. Amplification of 483 bp complete coding sequence of *gremlin* from hydra using PCR (a). Translated peptide sequence of Gremlin (b) and Noggin (c) shows N-terminal signal peptide (brown), conserved CTC domain (blue) and conserved Cysteine residues (red) involved in the formation of eight and ten membered ring respectively. Potential glycosylation sites, NXT/NXS in both protein sequences are shown in pink. Multiple alignment of gremlin (e) across different organisms (d) from hydra to vetebrates shows conserved C-terminal region. Sequence alignment between hydra Gremlin and Noggin shows variable N-terminal and conserved C-terminal regions (f).

### *In silico* analysis of Gremlin and comparison with Noggin

The translated peptide sequence of Gremlin from *H. vulgaris* Ind-Pune was analyzed using the protein database SMART which confirmed the presence of a characteristic C-terminal Cysteine Knot (CTCK/CT) domain or von Willebrand type C (VWC) domain (shown in blue), and a 25 amino acid residue secretory signal sequence at the N-terminus (shown in brown) (Fig. 1b). Cysteine residues participating in forming the 8-membered ring are shown in red. Sequence comparison by CLUSTALW analysis of Gremlin across different species (Fig. 1d) showed a variable N-terminal region and a more conserved C-terminal region (Fig. 1e). Analysis of hydra Noggin also revealed the presence of characteristic CTCK domain (shown in blue) in the C-terminus which is comprised of 9 Cysteine residues (shown in red) required for the formation of a 10-membered ring (Fig. 1c). Sequence alignment of different BMP antagonists among themselves showed variable N-terminal region with more similarity/identity towards the C-terminal region (Walsh et al., 2010). Similarly, alignment between hydra Gremlin and Noggin also revealed variable N-terminal region and increased similarity towards the C-terminal region (Fig. 1f). Both ligands are glycoproteins and show potential N-glycosylation sites on Asparagines (NXT/NXS), at 70^th^ and 101^th^ positions in Gremlin and at 141^st^ position in Noggin using NetNGly prediction tool. To understand the structural conservation of Gremlin and Noggin from Hydra to vertebrates, homology modeling was carried out for the available sequences from different animals. Predicted homology models generated using UCSF Chimera program based on the available crystal structure of human Gremlin-1 homodimer (pdb-5aej) and crystal structure of BMP-7-Noggin complex (1m4u.1) showed high structural/topological conservation of both proteins across phyla (Fig. 2). Ramachandran plot analysis of generated models using RAMPAGE application showed 100 % residues falling in favoured and allowed regions with 0 % outliers for hydra Gremlin and 97 % residues falling in favoured and allowed regions with 3 % residues in outlier region for hydra Noggin. Superimposed models of hydra Gremlin with counterparts of other species showed maximum root mean square (RMS) value with *Nematostella vectensis* (Fig. 2A), a fellow Cnidarian, while superimposed models of hydra Noggin with other species showed maximum deviation with *Ambystoma mexicanum* (Fig. 2B). Both models show less deviation with human counterparts suggesting their conservation from early metazoans to humans.

**Fig. 2.**
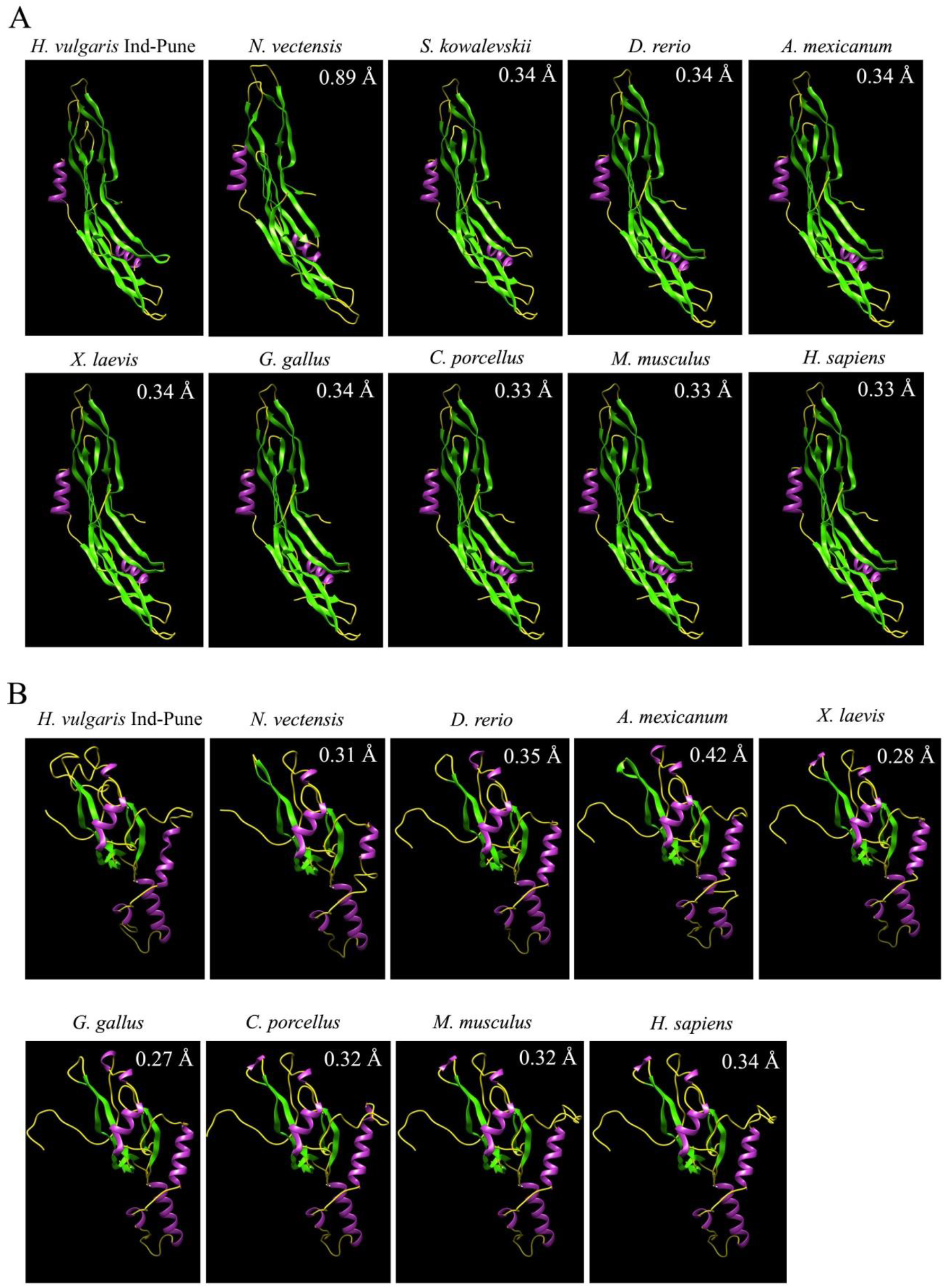
Homology models of hydra Gremlin and Noggin. Predicted homology models generated using UCSF Chimera program showed topological conservation of hydra Gremlin (A) and Noggin (B). Superimposed models of hydra Gremlin and Noggin shows maximum root mean square (RMS) deviation with *Nematostella vectensis* (0.89Å) and *Ambystoma mexicanum* (0.42 Å) respectively. RMS values for each organism are shown inset of each predicted model.

### Localization of *gremlin* and *noggin* in non-budding and budding polyps

Expression patterns of *gremlin* and *noggin* were assessed by whole mount *in situ* hybridization in whole polyps followed by serial transverse sectioning to study germ line specific localization. *Gremlin* transcripts were expressed ubiquitously in the endoderm of body column forming a gradient with highest expression in the budding region and lowest in the head and foot regions (Fig. 3Ab). The transcripts were predominantly expressed in the endoderm of budding region in whole polyps suggesting a possible role in budding. This was confirmed by *in situ* hybridization in polyps with different stages of buds, stage −3 (Fig. 3Ac), −4e (Fig. 3Ad), −5e (Fig. 3Ae) and −7 (Fig. 3Af), which showed predominant expression of *gremlin*, especially during the early stages, 3, 4e and 5e (arrows in Fig. 3Ac, d, e, respectively) when the new head organizer is established, confirming its role in budding. However, no expression was detected in the hypostome and foot regions that show organizer properties on transplantation (Browne, 1909; Gilbert, 2000; Kadu et al., 2012), suggesting the absence of direct involvement of *gremlin* in organizer formation in hydra. *Noggin* transcripts, on the other hand, showed significant expression in the endoderm of hypostome, base of the tentacles, lower body column and in the basal region in whole polyps (Fig. 3Bb). The observed expression of *noggin* in hypostome, base of the tentacles and foot region is comparable to our earlier observations made using *Xenopus noggin* probes in hydra (Chatterjee et al., 2001). During budding, *noggin* continued to be expressed in the hypostome, base of the tentacles, lower body column and basal disc in the adult polyp (Fig. 3Bc-f) and was also observed at the sites where new tentacles would emerge in the developing buds (shown by arrows in Fig. 3Bc, d, e). This suggests role of *noggin* in Organizer formation and tentacle formation. Expression of *noggin* in the hypostome and basal disc in whole polyps (Fig. 3Bb) that show Organizer activitities on transplantation (Browne, 1905; Gilbert, 2000; Kadu et al., 2012), points towards Organizer function of *noggin* in hydra.

**Fig. 3.**
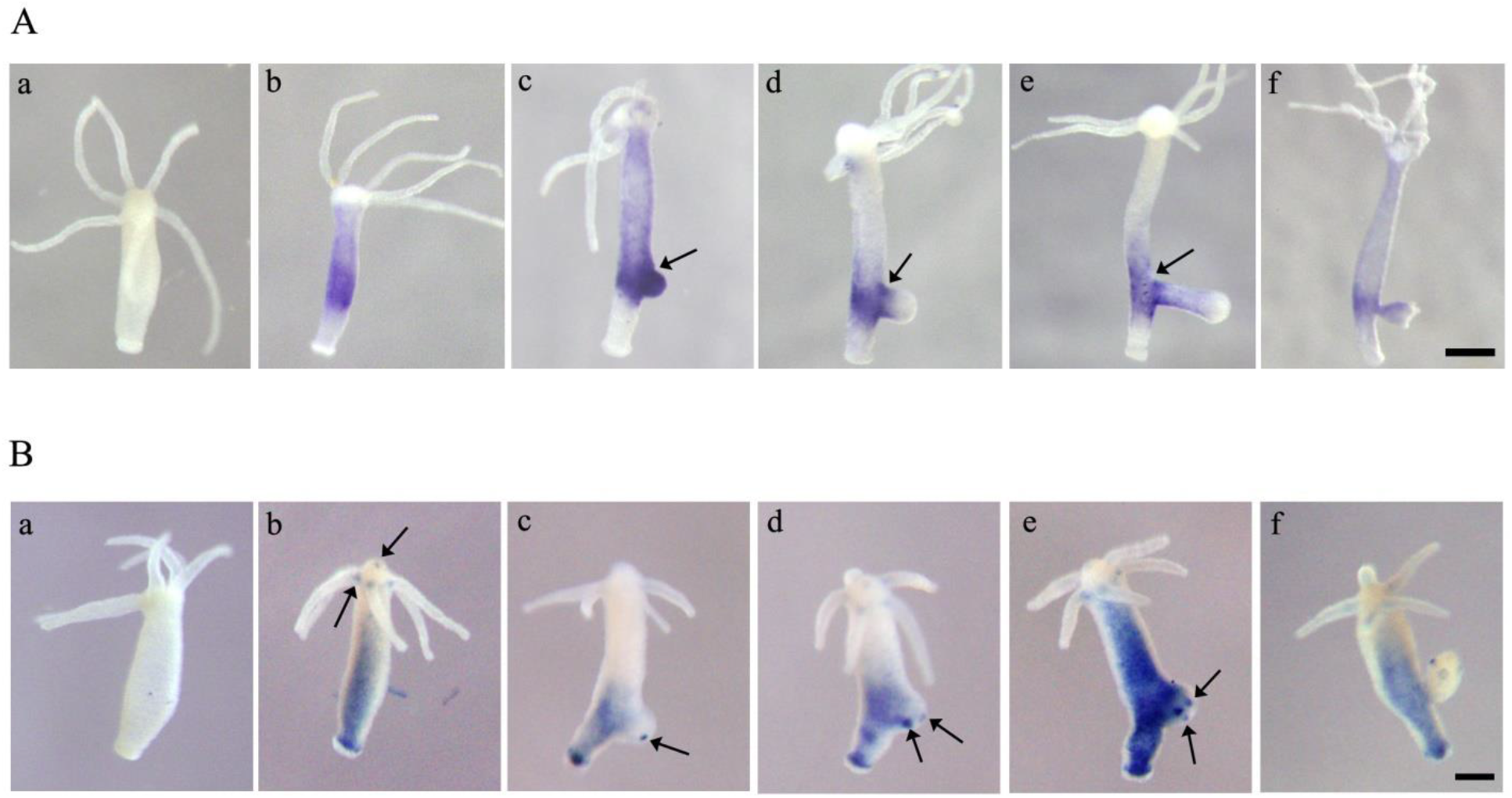
Localization of *gremlin* and *noggin* in hydra. Gremlin (A) is expressed in the budding region and body column of non-budding polyp (b), with no expression in the basal disc and hypostome. In developing buds, stages-3 (c), −4e (d), −5e (e) and −7 (f), predominant expression is seen during budding in the early stages (shown by arrows in Ac, d, e). *Noggin* expression (B) is predominant in the hypostome, base of the tentacles, lower body column and in the basal region in non-budding polyp (b). In developing buds, stages-3 (c), −4c (d), −5c (e) and −6 (f), *noggin* is seen as spots at the sites where new tentacles would emerge (shown by arrows in Bc, d, e). ‘Aa’ and ‘Ba’ represent polyps hybridized with sense probes for *gremlin* and *noggin*, respectively.

### Localization of *noggin* and *gremlin* with activated Wnt signaling

In view of the expression of *noggin* in the hypostome, base of the tentacles in whole polyps, and during bud development, its localization in alsterpaullone treated polyps was examined, since this experimental condition induces formation of ectopic Organizers and tentacles in the body column of the polyps which otherwise lacks Organizer property. This happens as a consequence of up-regulated Wnt signaling. For this, hydra treated with 5 μM Alsterpaullone for 24 h followed by recovery in hydra medium for different time intervals (48, 72 and 96 h) were used. Inactivation of GSK3β by alsterpaullone leads to over-activation of canonical Wnt pathway in hydra resulting in the formation of ectopic tentacles or multiple axes along the upper 2/3 portion of the body column. *In situ* hybridization showed expression of *noggin* as distinct spots where the ectopic tentacles would emerge after 48 h (Fig. 4Ad) post recovery in hydra medium. This spectacular expression of *noggin* persisted at the base of ectopic tentacles formed along the body column after 72 (Fig. 4Ae) and 96 h (Fig. 4Af) post recovery in hydra medium. This has been confirmed in transverse sections of *in situ* hybridized polyps which showed expression of *noggin* in the endoderm layer of body column (Fig. 4Bb) and foot (Fig. 4Bc) regions in whole polyps and at the base of ectopic tentacles in alsterpaullone treated polyps (Fig. 4Cb, c). Sections passing through polyps hybridized with sense probes show no signal in control (Fig. 4Ba) and alsterpaullone treated polyps (Fig. 4Ca). *In situ* hybridization in DMSO treated control hydra also showed significant expression of *noggin* in the hypostome, base of the tentacles and in the foot region at 48 (Fig. 4Aa,), 72 (Fig. 4Ab) and 96 h (Fig. 4Ac) post recovery in hydra medium. Expression of *noggin* in the body column as distinct spots in alsterpaullone treated polyps thus confirms the Organizer function of *noggin* in hydra. As opposed to *noggin* expression, *gremlin* expression showed diffused expression in the endoderm of body column in alsterpaullone treated polyps at 48, 72 and 96 h (Fig. 5Ab, c, and d, respectively). Magnified images (Fig. 5B) showed diffused expression in alsterpaullone treated polyps at 48 (Fig. 5Ba), 72 (Fig. 5Bb) and 96 h (Fig. 5Bc) post recovery in hydra medium with no expression in the original head (H) and foot (F) regions at all time intervals (Fig. 5Ba-c). Absence of *gremlin* transcripts in the hypostome (Fig. 5Ca) and foot (Fig. 5Cc) regions and in the body column (Fig. 5Cb) was also confirmed by transverse sectioning of control polyps. The finding that *noggin* is expressed in the hypostome and foot regions while *gremlin* is not points towards their distinct functions in hydra.

**Fig. 4.**
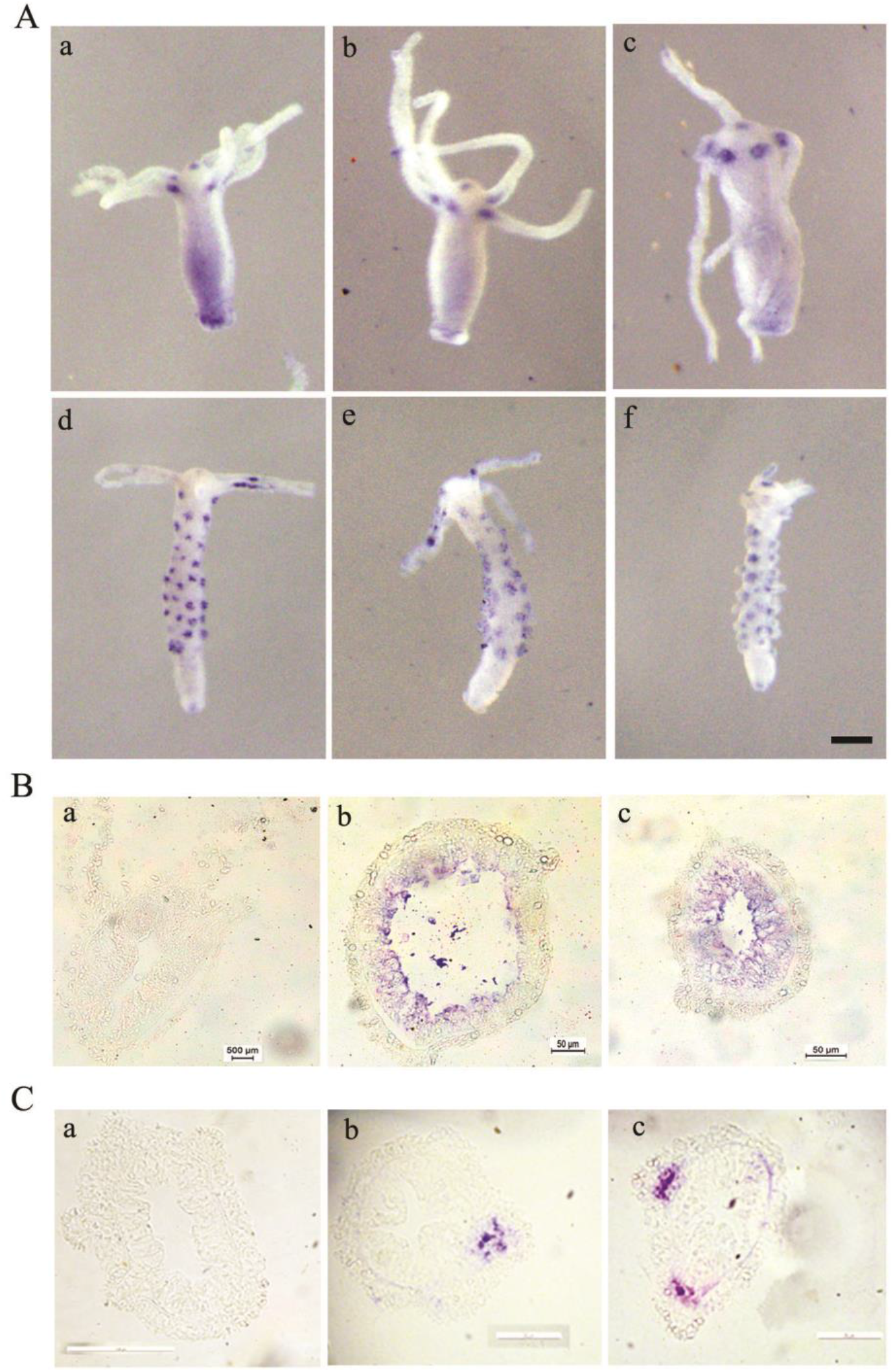
Localization of *noggin* in alsterpaullone treated polyps. Expression of *noggin* is observed as distinct spots in the 2/3^rd^ body column of alsterpaullone treated polyps (A) at 48 (d), 72 (e) and 96 h (f). *Noggin* expression in DMSO treated polyps is seen in hypostome, base of the tentacles and basal region (Aa, b, c). Scale bar = 200 μm. Sections passing through whole hydra (B) show endodermal expression of *noggin* in the body column (b) and foot region (c). Sections passing through alsterpaullone treated hydra (C) show endodermal expression of *noggin* at the base of the ectopic tentacles formed in the body column (b, c). ‘Ba’ and ‘Ca’ represent sections passing through the control polyp hybridized with sense probes.

**Fig. 5.**
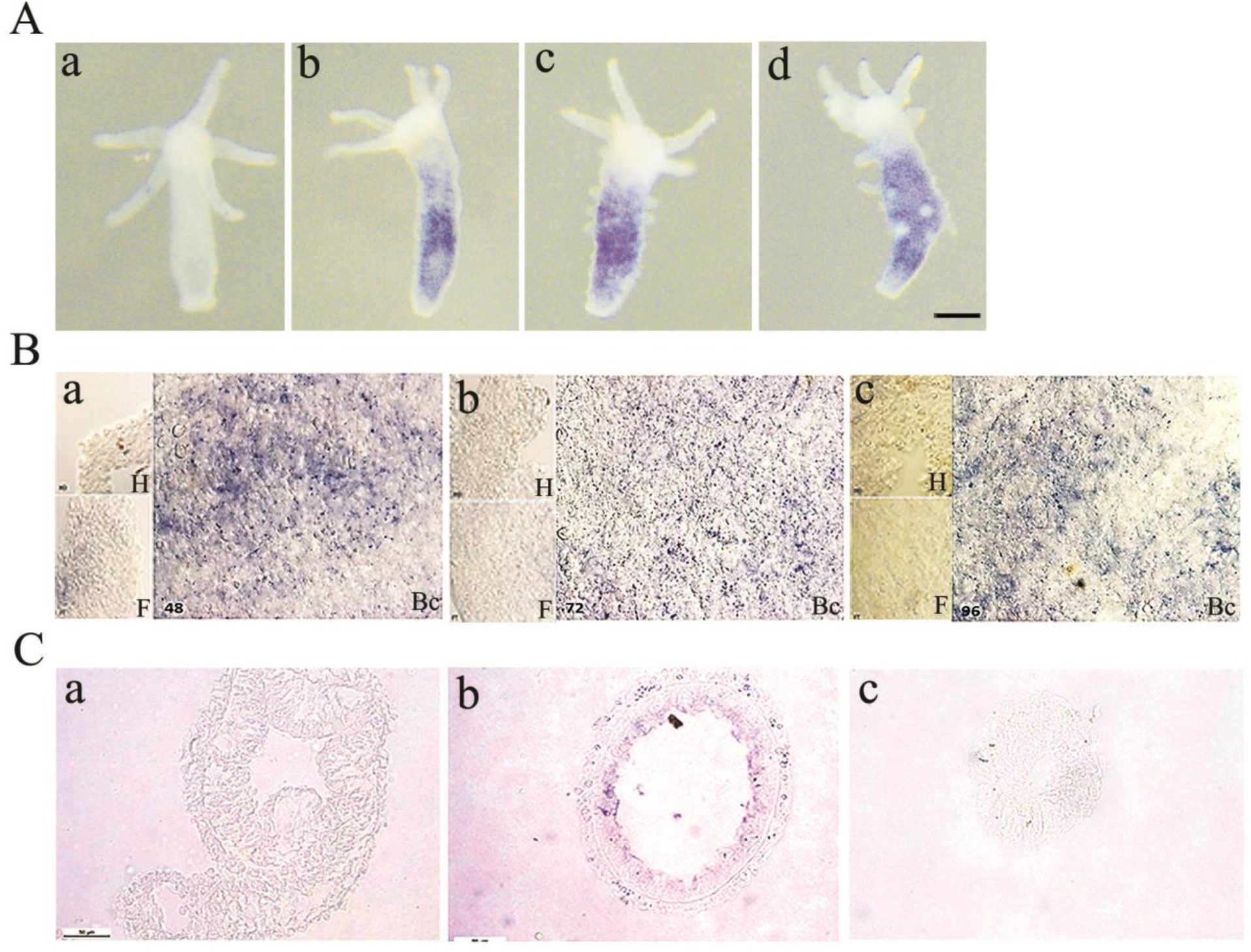
Localization of *gremlin* in alsterpaullone treated polyps. *Gremlin* (A) shows ubiquitous expression, except in the hypostome and foot region in alsterpaullone treated polyps (b, c, d). Scale bar = 200 μm. Magnified views of alsterpaullone treated polyps (B) show diffused expression in the body column (Bc) and show lack of expression in the head (H) and foot regions (F) after 48 (a), 72 (b) and 96 h (c). Sections passing through head, body column and foot regions of control polyp (C) show lack of expression in the hypostome (a) and foot (c) while endodermal expression is seen in the body column (b).

### Antagonism between Wnt and BMP pathways in hydra

Four major types of molecular interactions have been identified between BMP and Wnt pathways (Itasaki and Hoppler, 2009). To understand the nature of interaction occurring between these two pathways in hydra and to identify the possible involvement of *noggin* and *gremlin* during such interaction, canonical Wnt pathway was over-activated in the body column of hydra using alsterpaullone and expression levels of *BMP 5-8b, noggin* and *gremlin* were analysed by RT-PCR. Further, *FGF* levels were also checked, given that different types of interactions between FGF-BMP and FGF-Gremlin during tissue morphogenesis and limb bud formation have been reported (Verheyden and Sun, 2008; Zhu et al., 2014). *Wnt3* and *EF1-α* were used as positive and endogenous controls, respectively. Treatment with alsterpaullone for 24 h followed by return to hydra medium for 0.5 h (Fig. 6A) showed up-regulation of *Wnt3* by semi quantitative RT-PCR, confirming the activation of canonical Wnt signaling. Significant down-regulation of *BMP 5-8b* was observed as early as 0.5 h post-recovery in hydra medium which demonstrates the presence of antagonism between Wnt and BMP pathways. No significant change in the expression levels of *noggin* and *gremlin* was detectable at this time point (Fig. 6Aa). This suggests absence of direct involvement of both BMP inhibitors in causing down-regulation of *BMP 5-8b* by Wnt signaling in hydra. Also no change was detected in *FGF* levels suggesting lack of direct interaction between *Wnt* and *FGF* in down-regulating *BMP 5-8b*. Modulation of expression of *noggin*, *gremlin* and *FGF* was also examined after increased recovery time intervals in hydra medium. Recovery in hydra medium for 4 h also showed no significant difference in the expression levels of *noggin*, *gremlin* and *FGF*, while down-regulation of *BMP5-8b* still continued (Fig. 6Ba). However, at 24 h recovery, though *BMP5-8b* continued to get down-regulated, significant down-regulation of *FGF* was also observed for the first time, while no change was seen in expression of *noggin* and *gremlin* (Fig. 6Ca). At 48 h recovery, *BMP 5-8b* down-regulation continued, while relatively little down-regulation of *noggin* and *gremlin* was seen (Fig. 6Da). A careful observation of *BMP 5-8b* gene expression at all time intervals showed recovery of basal levels of *BMP5-8b* from 0.5 h till 48 h (Fig. 6Ea). Histograms plotted for the normalized values of *Wnt3, noggin, gremlin, BMP5-8b* and *FGF* against endogenous control, *EF1-α* at 0.5, 4, 24, 48 h showed no significant change in the expression levels of *noggin* and *gremlin* during *BMP5-8b* inhibition by Wnt pathway (Fig. 6 Ab; Bb; Cb; Db).

**Fig. 6.**
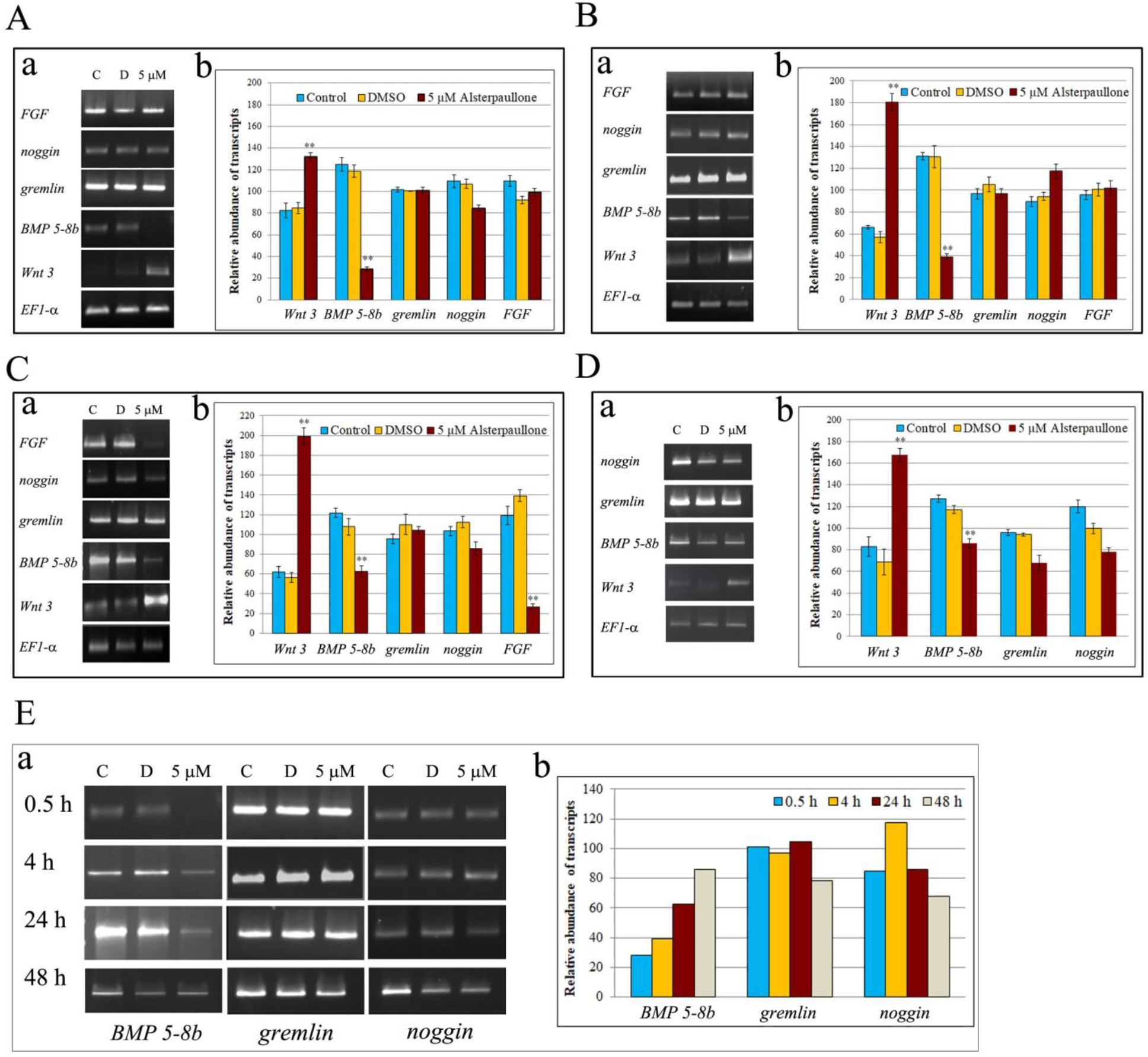
Effect of Wnt pathway on *BMP 5-8b* expression. Downregulation of *BMP 5-8b* at 0.5 h (Aa) post recovery in hydra medium was seen with activated Wnt signaling. Similar pattern of expression was seen at 4 (Ba), 24 (Ca) and 48 h (Da). No significant change was seen with *noggin* and *gremlin* expression at 0.5 (Aa), 4 (Ba) and 24 h (Ca). At 48 h post recovery, downregulation of both *noggin* and *gremlin* was seen (Da). No change in *FGF* levels was seen at 0.5 (Aa) and 4 h (Ba), while significant downregulation was seen at 24 h (Ca). Recovery of basal levels of *BMP5-8b* was seen from 0.5 h till 48 h, while no significant change was observed for *gremlin* and *noggin* (e). Histograms show normalized values of *Wnt3, noggin, gremlin, BMP5-8b* and *FGF* against *EF1-α* at 0.5, 4, 24 and 48 h (Ab; Bb; Cb; Db).

## Discussion

Establishment of anterior-posterior axis by canonical Wnt signaling is well documented in metazoans. Expression of *Wnt* in the posterior regions and development of anterior axis as a result of *Wnt* inhibition suggests that the anterior-posterior symmetry could be the ancestral condition in the body plan development of bilaterians (Petersen and Reddien, 2009). Similarly, in vertebrates, BMP signaling plays a crucial role in the patterning of dorso-ventral axis formation. A gradient of BMP signals, such as, high levels of BMPs specifies the ventral side, low levels specifies the lateral axis and lack of BMP signals leads to the dorsal side determination. Control of BMP signaling occurs by secreted glycoproteins such as Noggin, Chordin, Gremlin and Follistatin which bind to and inhibit BMP signaling. Likewise, role of FGFs in dorso-ventral axis specification in vertebrate embryos has also been demonstrated. It is thus clear that interactions between different signaling pathways are necessary in several developmental events including morphogenesis, cell fate specification, and organogenesis. Many of these pathways transduce signals by ligand binding to their respective receptors resulting in the activation of intermediate molecules, such as, secreted proteins and transcription factors that lead to the regulation of gene expression (Trompouki et al., 2011). Interactions between Wnt and BMP pathways have been well studied and involve distinct mechanisms. One of the important mechanisms by which these two pathways interact is through secretory molecules which bind to the extracellular components of Wnt and/or BMP pathways. In the present study, we have studied various aspects of BMP and Wnt signaling in hydra, a diploblastic organism with a simple but definite body plan that is believed to have evolved over 600 million years ago. We have characterized *gremlin* from hydra, compared its gene expression pattern with another BMP inhibitor, *noggin* and deduced their respective functions and their relationship with Wnt signaling in hydra.

Gremlin belongs to the CAN (Cerberus and DAN) subfamily of BMP antagonists that contains a C-terminal Cystine knot with 8-membered ring, resembling the ring structure of BMPs (Avsian-Kretchmer and Hsueh, 2004). The typical structure of 8-membered ring in these members contains Cysteine residues arranged in the form of C-X_n_-CXGXC-X-C_1/2_-X_n_-CXC (Avsian-Kretchmer and Hsueh, 2004). *In silico* structural analysis of hydra Gremlin also revealed conservation of the N-terminal signal peptide and a characteristic CTCK domain in the C-terminus. The motif in hydra Gremlin shows C-X_9_-CXGXC-X_22_-CX_2_C-X_13_-C-X_17_-CXC arrangement, where ‘C’ represents Cysteine residue involved in forming the ring, ‘X’ represents any amino acid and numbers in the subscript represent number of amino acids present between two Cysteine residues. This suggests structural conservation of Gremlin from hydra to humans. This is further validated by generating homology models for hydra Gremlin using the crystal structure of human Gremlin, which showed less RMS deviation upon superimposition with human Gremlin. A similar observation is made for hydra Noggin which showed a characteristic 10-membered ring motif CX_22_CX_5_CX_7_CX_14_CX_10_CX_12_-CXCXC as seen in human Noggin (Avsian-Kretchmer and Hsueh, 2004). Hydra homology models generated based on the human BMP-Noggin complex crystal structure also showed less deviation with human Noggin. This suggests the structural conservation of both BMP inhibitors from hydra to vertebrates including humans.

Localization studies by whole mount *in situ* hybridization followed by serial transverse sectioning revealed *gremlin* expression predominantly in the endoderm of budding region and early stages of bud formation in hydra. This suggests that *gremlin* is possibly involved in budding in hydra. It is established that Gremlin is the principal BMP inhibitor that plays a key role in limb bud development in vertebrate embryos (Verheyden and Sun, 2008). Also it is reported that *gremlin* regulates *FGF* and *sonic hedgehog* to direct the outgrowth of the limb (Khokha et al. 2003). Limb bud formation in vertebrates and bud formation in hydra involve accumulation of cells resulting in the formation of tissue evagination and protrusion forming a circular bulge. Expression of *gremlin* in the budding region (whole polyps), during bud evagination (stage 3-4 buds) and protrusion (stage 5-6 buds) in hydra suggests presence of similar mechanisms during budding (see Fig. 3Ac, d). Previous studies have reported expression of *gremlin* in the migrating neural crest cells and not in the Organizer, suggesting its potential role in the overall development of embryo and not restricted to gastrulation (Hsu et al. 1998). Similarly, we do not detect expression of *gremlin* in the hypostome and foot that show Organizer function in hydra (Browne, 1905; Gilbert, 2000; Kadu et al. 2012) indicating that *gremlin* is not involved in Organizer function in hydra. Lack of *gremlin* expression in the regenerating tips (Fig. S1) further supports this conclusion.

*Noggin* transcripts, unlike *gremlin* transcripts, showed significant expression in the endoderm of hypostome, base of the tentacles, lower body column and in the basal region in whole polyp and at sites where new tentacles would emerge in the developing bud. This suggests role of *noggin* in tentacle formation. Expression of *noggin* in the hypostome and basal disc in whole polyps that show Organizer activity (Browne, 1905; Gilbert, 2000; Kadu et al., 2012) points towards the role of *noggin* in Organizer formation in hydra. This is further confirmed in polyps with activated Wnt signaling which showed *noggin* expression as distinct spots in the body column where the tentacles would emerge from newly formed ectopic Organizers (see Fig. 4). It is interesting to note that this expression pattern of *noggin* as distinct spots before the emergence of tentacles in the developing buds and in the alsterpaullone treated polyps exactly coincides with the reported expression pattern of *BMP5-8b* in hydra (Reinhardt et al. 2004). The first tentacles formed in the developing bud face the foot of the parent polyp and show *BMP 5-8b* expression as distinct spots in the young bud (Reinhardt et al. 2004). *Noggin* expression appears in the same region as two spots facing the foot of the parent polyp in the developing bud (Fig. 3Bd), suggesting the presence of interaction between *noggin* and *BMP5-8b* during tentacle patterning and morphogenesis. In alsterpaullone treated polyps, both *noggin* and *BMP5-8b* showed similar expression pattern as spots on the body column before the emergence of tentacles and at the tentacle base zone once the tentacles are formed. This perfectly overlapping spatiotemporal expression of *noggin* and *BMP5-8b* in suggests their role in displacing the tissue from the body column to tentacle border zone and then to tentacles. Thus, *noggin* and *BMP5-8b* seem to be involved in either tentacle patterning, morphogenesis or in both. However, we could not detect expression of *noggin* at the regenerating ends of head and foot pieces (Fig. S2).

The findings that Wnt signaling modulates BMP pathway in different ways prompted us to study the expression of *BMP5-8b*, *gremlin* and *noggin* with activated Wnt signaling in hydra. Since FGF signaling plays a crucial role in the establishment of anterior-posterior axis patterning during embryo development (Dorey and Amaya, 2010) and interactions between FGF-BMP and FGF-Gremlin during tissue morphogenesis and limb bud formation have also been reported (Verheyden and Sun, 2008; Zhu et al., 2014), transcript levels of *FGF* were also analyzed. Though Wnt and BMP pathways can and do act independently, both co-operative and antagonistic mechanisms exist between them depending on the cell and tissue type. Four types of functional and molecular interactions between them have been identified depending on the cellular context and it is important to understand such interactions during development (Itasaki and Hoppler, 2010). In hydra, activated Wnt signaling resulted in localization of both *noggin* and *BMP5-8b* as distinct spots on the body column. In order to identify the nature of interactions between *Wnt* and *BMP5-8b* in hydra, semi-quantitative RT-PCRs were performed for *BMP5-8b* in alsterpaullone treated hydra after different time intervals post transfer to hydra medium. Treatment for 24 h followed by return to hydra medium for 0.5 h showed up-regulation of *Wnt3*, confirming the activation of canonical Wnt signaling. We find significant down-regulation of *BMP 5-8b* with activated Wnt signaling demonstrating the presence of antagonism between these pathways in hydra. Though molecular interactions (either synergism or antagonism) between Wnt and BMP pathways have been well studied in different cellular contexts, inhibition of BMP pathway by Wnt signaling has been reported only under few conditions. For example, in *Drosophila* leg development and in eye/antennal discs, wingless (wg) and decapentaplegic (Dpp), a *Drosophila* homolog of human BMP2/4, show antagonistic relationship by repressing each other’s expression thus providing distinct territories for both during development (Theisen et al., 1996). Also induction of neural tissue by Wnt8 by inhibition of *BMP4* expression is reported in *Xenopus* embryos (Baker et al. 1999). In other cases, expression of Wnt may induce secretory molecules that bind to and inhibit BMPs such as PRDC, Xiro1, BMP-activin membrane bound inhibitor, thereby resulting in inhibition of BMP pathway (Glavic et al., 2001; Sekiya et al., 2004; Im et al., 2007). Up-regulation of *gremlin* by *Wnt* in human fibroblasts (Klapholz-Brown et al., 2007) and inhibition of *BMP4* by inducing Noggin as a result of *Wnt1* activation during somite patterning in chick embryos has also been reported (Hirsinger et al., 1997). Here, we investigated whether up-regulation of *noggin* and/or *gremlin* by *Wnt* has resulted in *BMP 5-8b* down-regulation. This was not so as RT-PCR results showed no significant change in the expression levels of *noggin* and *gremlin*. Thus, these two BMP inhibitors are not involved in down-regulation of *BMP 5-8b* by Wnt signaling in hydra. Also, no change was detected in *FGF* levels in the early time points (0.5 and 4 h) suggesting the absence of direct interaction between *Wnt* and *FGF* in down-regulating *BMP 5-8b*. Significant down-regulation of *FGF* was seen only after 24 h. It is known that FGFs act as posteriorising factors (Dorey and Amaya, 201). With activated Wnt signaling in hydra, the body column takes head fate resulting in the formation of ectopic tentacles and organizers after 24 h post transfer to hydra medium. Down-regulation of *FGF* at this time interval thus confirms the presence of interactions between *Wnt3* and *FGF* in hydra. It is interesting to note that with increased recovery time intervals in hydra medium, expression of *BMP5-8b* increases. Slight down-regulation of *noggin* and *gremlin* was also seen at 48 h. It is known that low levels of BMPs cause up-regulation of *noggin* and *gremlin*, while high concentrations of BMPs inhibit *noggin* and *gremlin* (Re’em-Kalma et al., 1995; Gazzero et al., 1998; Nissim et al., 2006). The observed down-regulation of *noggin* and *gremlin* at 48 h recovery, therefore, could be the result of constant expression of *BMP5-8b* at the base of the tentacle.

In summary, in addition to identification and *in silico* characterization of hydra Gremlin, our results show differential expression of BMP inhibitors *gremlin* and *noggin* in hydra and demonstrate the absence of direct involvement of *gremlin* and *noggin* in inhibition of BMP pathway by Wnt signaling in hydra. Most importantly, our data indicate that BMP/Noggin antagonism as a mechanism for Organizer formation is evolutionarily ancient. Gremlin and Noggin may have been recruited for different/additional functions during vertebrate axial patterning. A better understanding of the roles of BMP antagonists and the interplay between various pathways in hydra, which lacks a dorso-ventral axis, would help in understanding the evolution of body axes and body plans in metazoans.

## Materials and Methods

### Hydra culture

Clonal cultures of *Hydra vulgaris* Ind-Pune (Reddy *et al*., 2011) were maintained in hydra medium (Sugiyama and Fujisawa, 1977) in glass crystallizing bowls at a constant temperature of 18 ± 2°C with 12 h light/dark cycle. Polyps were fed with freshly hatched *Artemia salina* nauplii on alternate days.

### *In silico* analysis

Sequence alignments for peptide sequences of Gremlin were carried out using CLUSTALW analysis. Homology models for Gremlin and Noggin were generated in UCSF Chimera based on the pdb files generated by both manual and automated methods using Swiss model work space and compared with homology models generated for available sequences of different organisms across phyla.

### Whole mount *in situ* hybridization

Whole-mount *in situ* hybridization using digoxygenin (DIG)-labeled RNA probes was carried out as previously described (Krishnapati and Ghaskadbi, 2013) with few modifications. Briefly, pGEMT Easy vector harboring complete coding sequences of *gremlin* and *noggin* clones were amplified using T7 and SP6 promoter primers, purified and used for *in vitro* transcription reaction using Dig-RNA labeling kits (Roche). Following *in situ* hybridization, serial transverse sectioning of hydra was performed to study details of expression patterns as previously described (Krishnapati and Ghaskadbi, 2013).

### Primer design, PCR and statistical analysis

Sequences flanking the open reading frame of *gremlin* and *noggin* mRNAs were used to design primers for amplifying the complete coding sequences. Analysis of expression of desired genes was carried out by semi quantitative RT-PCR using cDNA as template. Each experiment was carried out at least in triplicate. Histograms were computed by normalizing the values of band intensities of test genes with *Hyactin/EF1-a*. Mean and standard deviation were calculated for each experimental set and statistical significance was calculated by Students paired ttest.

### Treatment with alsterpaullone

Hydra starved for 24 h were treated with 5 μM alsterpaullone, an inhibitor of GSK-3β, for 24 h, as described previously (Broun et al., 2005). Subsequently, hydra were thoroughly washed with hydra medium and transferred to fresh medium for different time intervals, viz., 0.5, 4, 24, 48, 72 and 96 h. Hydra polyps treated with appropriate concentrations of dimethyl sulfoxide (DMSO) served as solvent controls while those in hydra medium served as master controls.

## Acknowledgements

We thank Dr. Vidya Patwardhan and Ms. Rohini Londhe for discussions and help and Ms. Aditi Kavimandan for help in whole mount *in situ* hybridization.

---

**S1.**
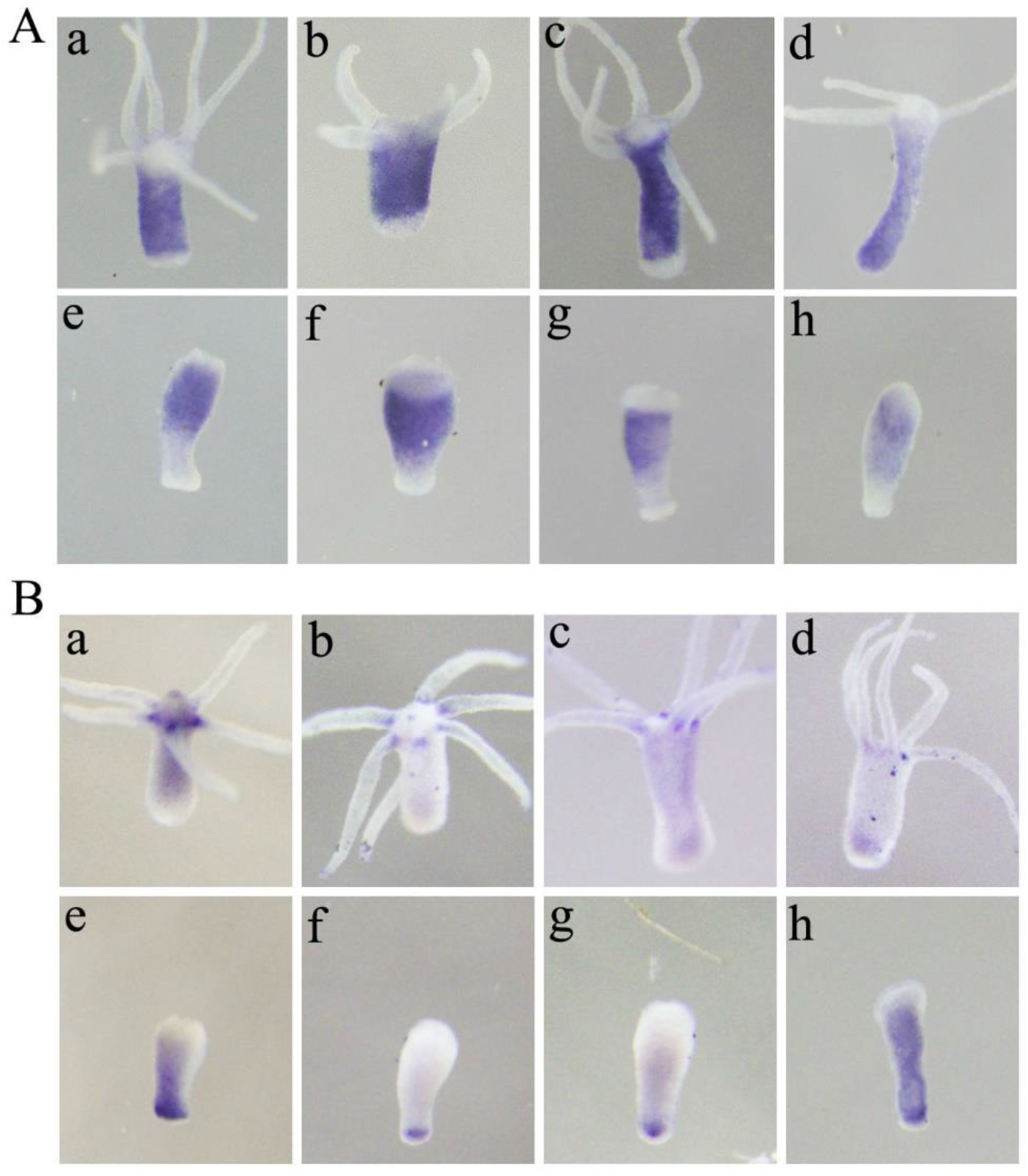
*Gremlin* and *noggin* expression in regenerating hydra. No expression of *gremlin* (A) was seen in foot regenerating (a-d) and head regenerating (e-h) pieces after 1, 2, 4 and 24 h respectively post mid-gastric bisection. No significant expression of *noggin* (B) was seen in both foot regenerating (a-d) and head regenerating (e-h) pieces at 0, 2, 4 and 18 h post mid-gastric bisection. Original expression of *gremlin* (A) in the budding region of foot pieces and *noggin* (B) in the hypostome, base of the tentacles in head pieces and at the basal disc in foot pieces is observed.

